# Water plant functional groups: a trait-based classification database for Australian plants

**DOI:** 10.64898/2026.10.18.759887

**Authors:** David C. Deane, Sharon Bowen, Cherie J. Campbell, Fiona J. Dyer, Fiona L Freestone, Caitlin V. Johns, Joanne E. Ling, Tanya J. Mason, Kaylene Morris, Jason M. Nicol, Alica Tschierschke, Lyndsey M. Vivian, Michelle T. Casanova

**Author notes:** ***Corresponding author:*** D.C. Deane Address: Dept. of Ecological, Plant and Animal Sciences, La Trobe University, Melbourne, Australia. ***Disclosure statement***: the authors have nothing to disclose.

## Abstract

Trait-based functional classifications of species offer improved generality in predicting responses to environmental conditions. Water plant functional groups (WPFG) are one such classification. Grouping plant species based on traits associated with germination, growth and reproductive requirements, WPFG have proven useful in examining plant trait-hydrological relationships in aquatic plant research in Australia. Here, we collate existing information from across academic, government and personal databases to create the most comprehensive compilation of plant taxa assignments to WPFG available. Taxonomic nomenclature was first standardized using accepted names under the Australian Plant Census/Australian Plant Name Index. We then assessed agreement among source classifications and derived a consensus-weighted fuzzy classification that retains uncertainty in WPFG assignment at both the species- and genus-level. The resulting database provides evidence-based classifications for the most commonly observed water plants in temperate Australia, including evidence of affinity to all functional groups. This approach supports greater consistency, transparency and reproducibility in WPFG assignment across studies while allowing users to select the level of classification most appropriate to their application. The database is freely available for download from the Zenodo digital repository.

## Introduction

Ecologists have long recognised the value of grouping animal species exploiting common resources or habitats into guilds for understanding patterns in community structure or coexistence (e.g, foliage gleaning birds; Root 1967). However, as plants share similar resource needs, they are more often grouped based on shared functional traits that determine their ability to respond to stressful, disturbed or competitive environments (Grime 1979, Noble and Slatyer 1980). Adaptation to flooding-induced anoxia is a major driver of wetland vegetation zonation, although other environmental factors, including salinity, can also influence vegetation structure and composition (Weiher and Keddy 1995).

The use of functional groups in aquatic plant communities can be traced back at least as far as van der Valk (1981), who defined twelve possible groups through combination of three life history traits (life span, propagule longevity, propagule establishment requirements). The functional group approach for aquatic plants acquired greater prominence during the 1990s with foundational studies in Europe (Hills et al. 1994), North America (Boutin and Keddy 1993) and Australia (Brock and Casanova 1997). The Australian study introduced the Water Plant Functional Group (WPFG) classification, which has since been widely applied to questions relating to riverine and wetland habitat responses to water regime (see ‘Prior uses of WPFG in research’). The WPFG framework has been highly influential, with the two main source publications (Brock and Casanova 1997, Casanova and Brock 2000) cited more than 1400 times (Google Scholar, accessed 30 June 2026).

Most prior applications of the WPFG classification have been motivated by the need to quantify environmental water requirements in a consistent manner over the large spatial scale associated with the Murray-Darling Basin (Fig.1). The framework is also relevant to environmental-water planning, wetland assessment and monitoring, delineation and restoration decisions that require consistent interpretation of vegetation responses across sites and regions (Campbell et al. 2014). Despite its widespread use, species assignments to WPFGs are scattered across disparate sources, including primary and technical literature and the personal databases of expert aquatic botanists. This fragmentation limits access to species assignments and hinders assessment of consistency and reproducibility between studies. Here, we describe a project that addresses this issue through the development of a database of prior assignments of species to WPFG. The collated database is unprecedented in (i) providing a consolidated and accessible repository of multiple lists assigning plants to a WPFG classification, (ii) quantifying variation in assignments among observers and study systems, and (iii) providing a fuzzy classification framework that captures alternative WPFG assignments and enables researchers to select the classification most appropriate to their study context. The database is freely available via the Zenodo digital repository at URL https://doi.org/10.5281/zenodo.23128087.

**Figure 1.**
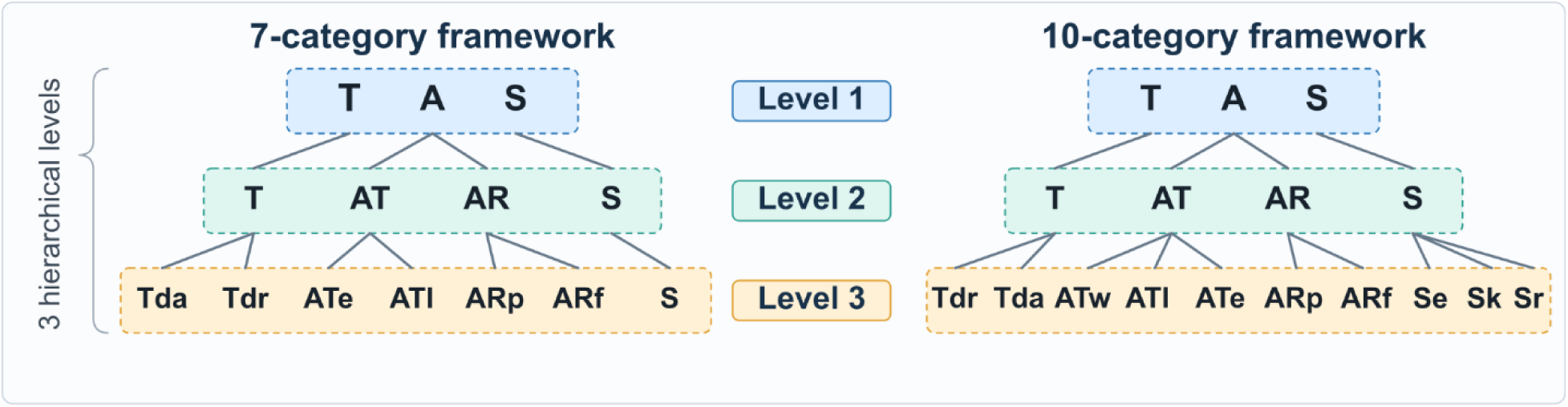
Alternative WPFG frameworks. Dashed lines enclose different categories at each hierarchical level.

### The origins and development of the Water Plant Functional Group classification

The origins of the Australian water plant functional group classification system lie in experimental work to understand wetland plant responses to wetting-drying cycles in temporary wetlands of the New England Tableland in the Australian state of New South Wales (Brock and Casanova 1997). This work combined field observations with germination and related experimental trials to examine plant performance under the highly variable wetting–drying regimes characteristic of Australian wetland habitats. Brock and Casanova (1997) and Casanova and Brock (2000) derived the classification by comparing germination, growth, and reproductive outcomes from seed bank samples of temporary wetlands under experimental variation in depth and duration of inundation. Through discriminant analysis, species responses were classified hierarchically into three, four or seven categories, which were then inductively linked to shared group-level traits.

The initial division into three classes is purely hydrological, splitting species into terrestrial (analogous to ‘upland’ in North American wetland indicator species classifications), amphibious, and submerged classes (designated T, A, and S), reflecting the amount and variability of ‘free water’ where established plants were typically found. The second level split in the hierarchy identifies two functional responses in amphibious species, differentiating: (i) tolerators – those species coping with the absence, presence or fluctuation of water levels without changing their morphology, from (ii) responders – species that actively alter their growth pattern (e.g., root-shoot ratio, leaf morphology), yielding a four-category classification (T, AT, AR, and S). Notably, the division of aquatic species into those that tolerate vs those that adapt to flooding has been independently identified from contrasting physiological adaptations to low oxygen availability (Bailey-Serres and Voesenek 2008), and in meta-analysis of wetland plant functional trait responses (Li et al. 2025), placing this division on sound theoretical ground. However, WPFG primarily distinguishes responders using visible morphological plasticity or elongation of leaves and stems; physiological or internal morphological adaptations are not necessarily separated by the classification. A similar split at the second hierarchical level differentiates the terrestrial group into those preferring generally drier (Tdr) conditions, which are widespread in disturbed terrestrial systems generally, from those preferring damp soil conditions (Tda), which are generally restricted to wetland habitats or moister microclimatic settings.

The third level in the Brock and Casanova (1997) hierarchy further differentiates the amphibious groups, separating tolerators based on growth habitat into emergent (ATe; serotinous seed dispersal rather than dependence on the seed bank) and low-growing (ATl; reliant on wind-dispersal rather than the seed bank) groups, and responders with plasticity of growth form (ARp) from free-floating plants or those with floating leaves (ARf). This results in the original seven-category classification (Tdr, Tda, ATl, ATe, ARp, ARf, S) introduced in Brock and Casanova (1997), with later work (Casanova and Brock 2000), further elucidating the role of water regime. A subsequent study of environmental water requirements in an intermittent stream catchment in South Australia led Casanova (2011) to propose an additional amphibious tolerator group based on woody growth form and serotinous seed dispersal rather than dependence on the seed bank (ATw) and to split the submerged group into three subclasses: emergent and reliant on wind-dispersal rather than the seed bank (Se) and, using r/K selection theory, Sr and Sk. This resulted in the full 10-category WPFG classification.

Many studies have successfully applied adaptations of the WPFG classification, ranging from the most general three-category (T, A, S) structure to the full 10-category classification proposed by Casanova (2011) (Table 1). The 10-category classification has been considered overly specific to the study system where it was developed for general use (Catford et al. 2017), but it is not uncommon for researchers to include additional categories. For example, under the Flow-Monitoring, Evaluation and Research (Flow-MER) program informing adaptive management in the Murray–Darling Basin (Dyer et al. 2021, Dyer et al. 2025; Table 1), woody species (ATw) are split into three categories (flow dependent, woody, other) to best match reporting requirements. The hierarchical nature of the classification makes it straightforward to add or reduce complexity and the number of categories adopted is best viewed as a design decision that should be informed by the study aims.

**Table 1.**
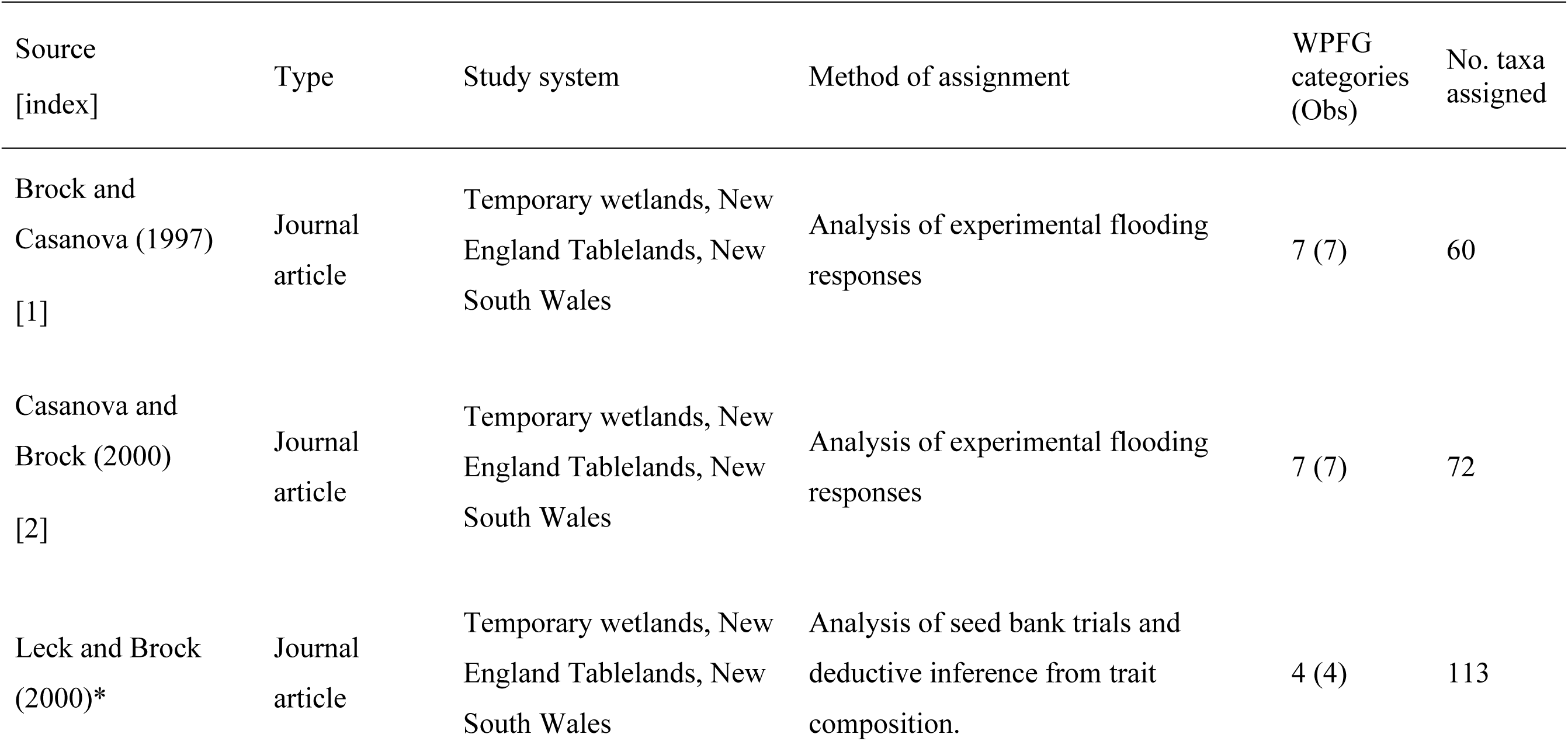
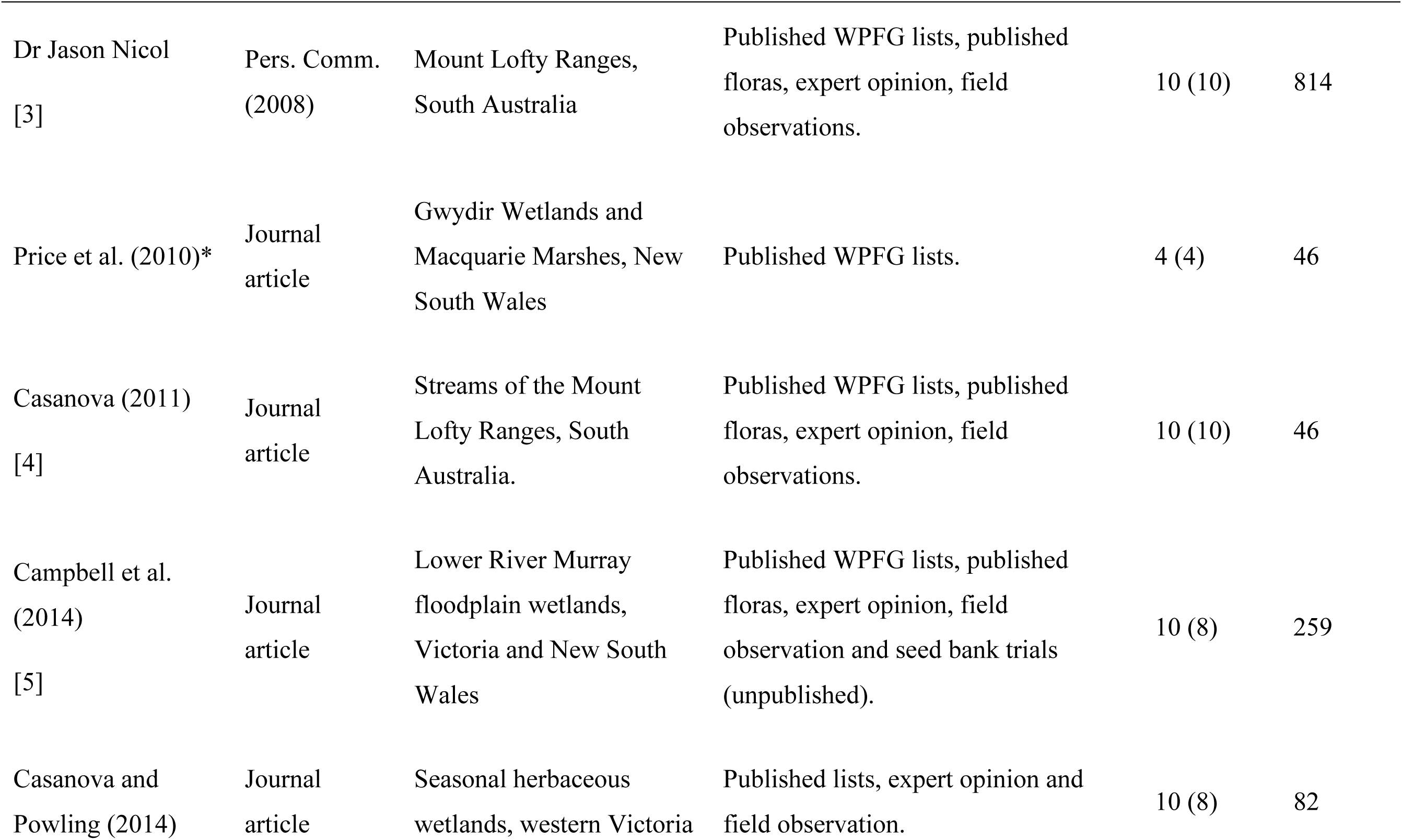
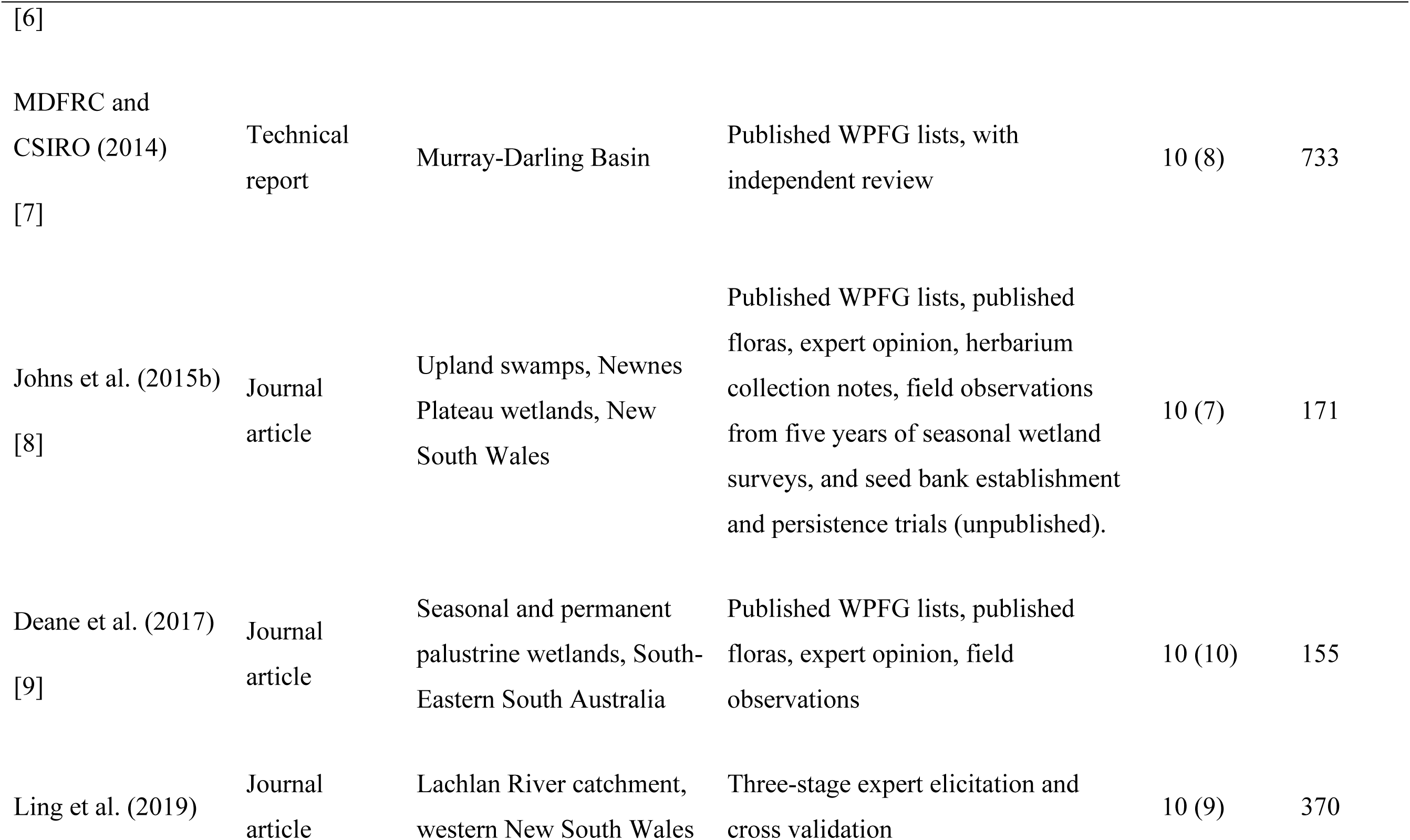
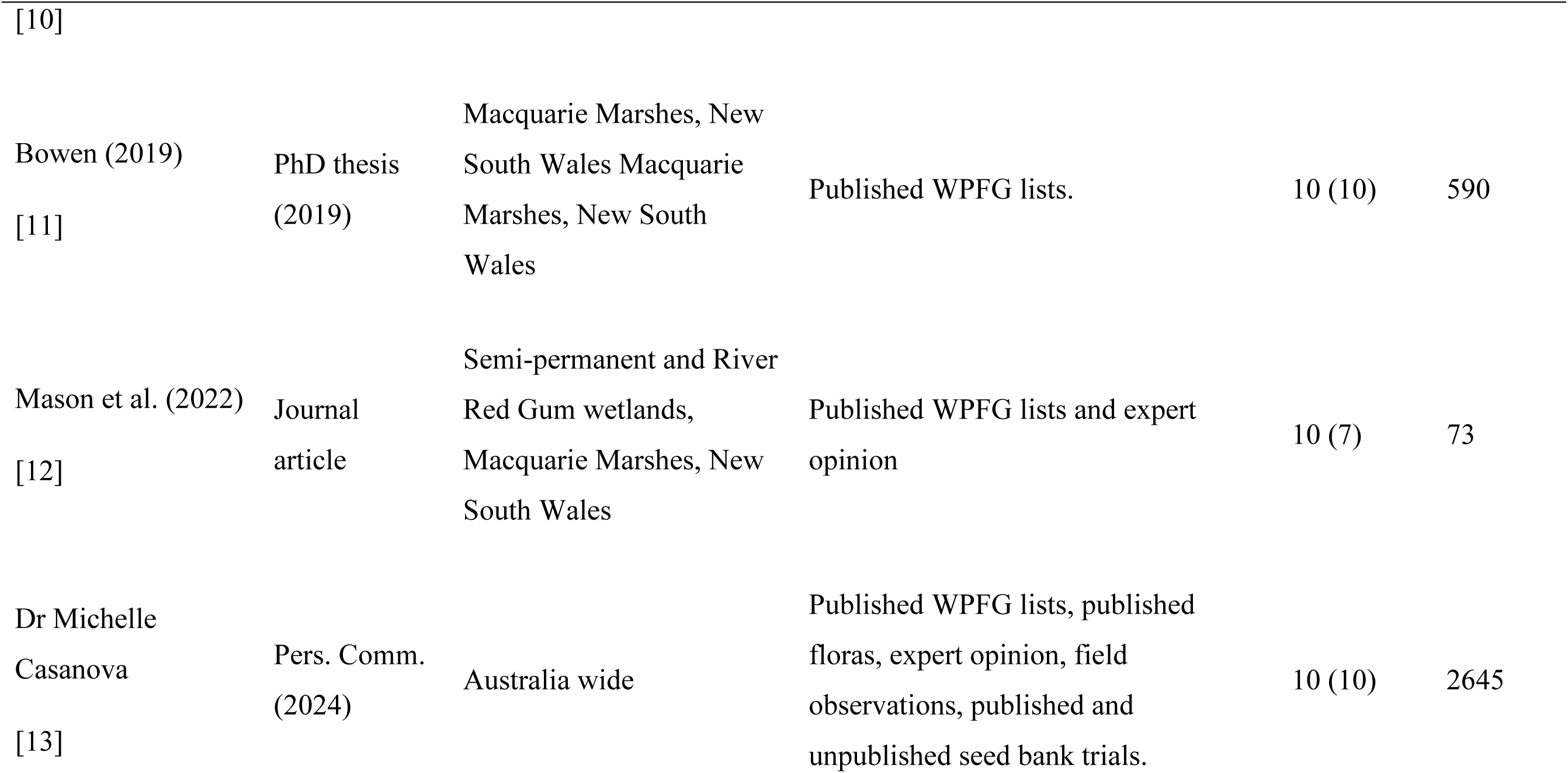
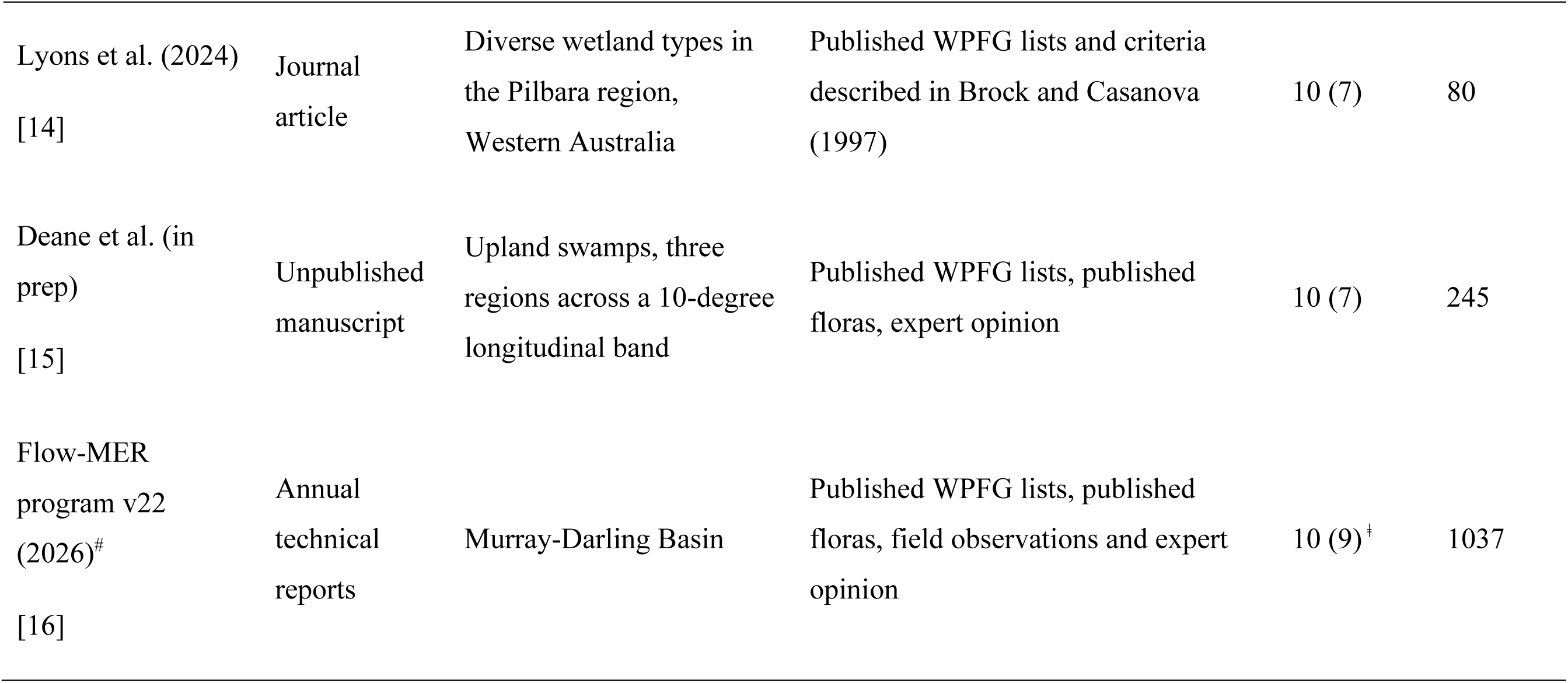
Prior studies in the development or assignment of species to water plant functional group categories with associated metadata. ‘WPFG categories’ refers to the number of alternative categories into which species were classified. Because not all allowed categories are present in every list, the actual number observed is also shown (Obs). Only studies assigning to the original 7 or the revised 10 category classification are included in the database (*denotes studies included in this table for historical context but not used in database) and included lists have an integer code [index] which gives the chronological order in which they were combined. ^#^Flow-monitoring Evaluation and Reporting project database as of 2026. Flow-MER 2019-2026 builds on the Long-Term Intervention Monitoring 2014-2019, producing annual technical reports. ǂ ignores three woody groups, pooling these into ATw for database inclusion.

### Recommended terminology

Given the different classifications in common use, we suggest describing Water Plant Functional Groups as a *framework* with hierarchical *levels* and differing numbers of *categories* at each different level (Fig. 1). For clarity, researchers should refer to the WPFG framework and specify the number of categories used (e.g., the 7-category or 10-category classification at the lowest hierarchical level), along with any additional modifications made. The assignment of taxa to categories is their *classification* within the WPFG framework (i.e., “Species X = Tda” is an example classification of a taxon to a category).

## Results

### Number of species and distribution among WPFG

Geographically, studies were heavily concentrated in the south-east of the country, especially the Murray-Darling Basin (Fig. 2). From the 16 contributed lists (Table 1, Fig. 2), a total of 6916 unique taxonomic assignments to WPFG were recorded, resulting in 3062 assignments to species level (or below) of which 3055 (99.7%) matched accepted nomenclature against the Australian Plant Census. Taxonomic richness declined strongly across categories associated with increasing adaptation to more persistent saturation and flooding (Fig. 3). Each unique taxon was assigned a best guess and associated fuzzy classification for all categories in both the seven- and ten-category systems. Within-genus functional-group composition and fuzzy classifications were analysed for 854 unique genera.

**Figure 2.**
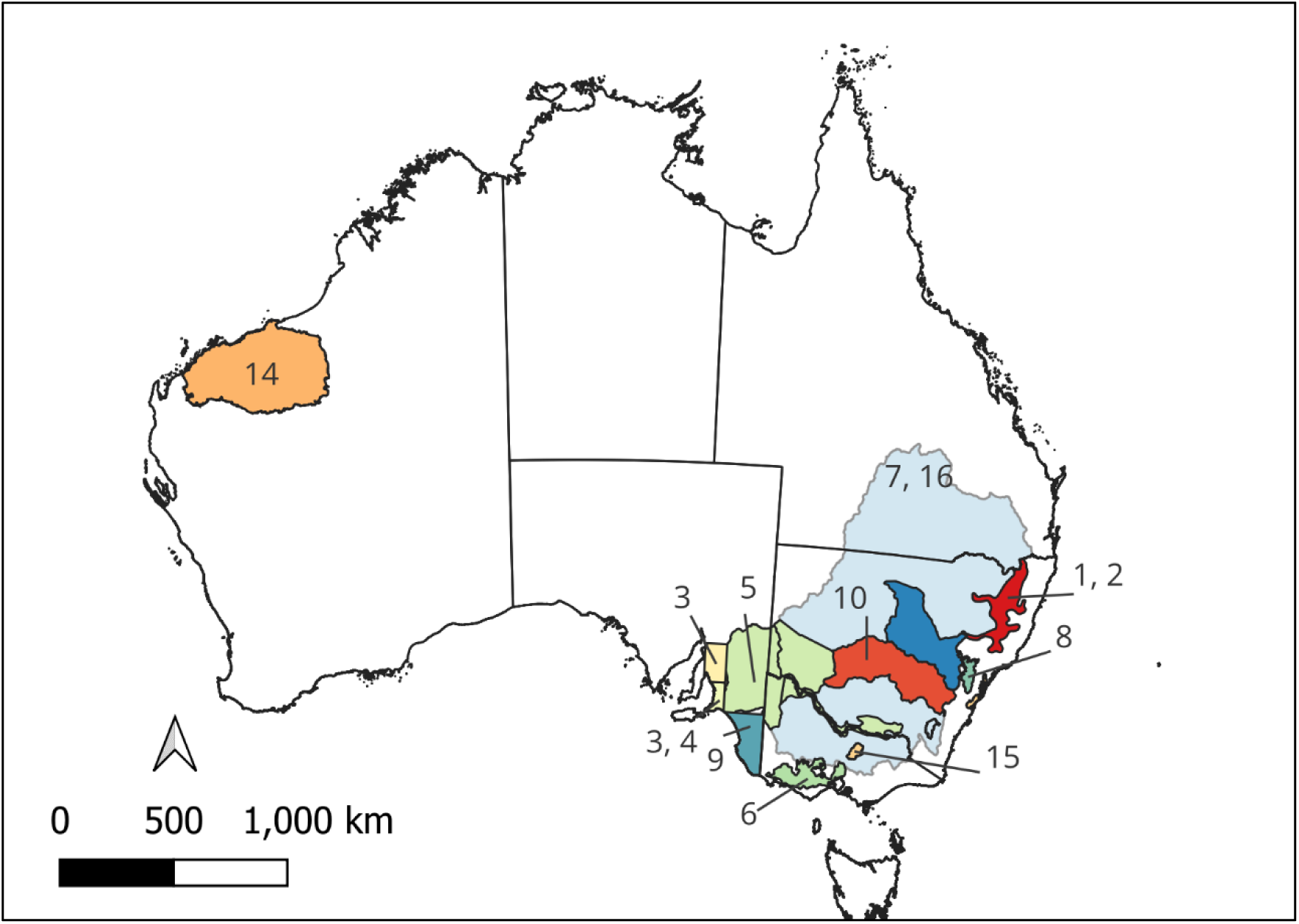
Approximate extent of study systems associated with each WPFG source listing collated within the database. Integers refer to study indices in Table 1. Source 13 can be thought of as a general nationwide listing, without specific geographical focus but is likely most appropriate in areas where prior WPFG studies have occurred.

**Figure 3.**
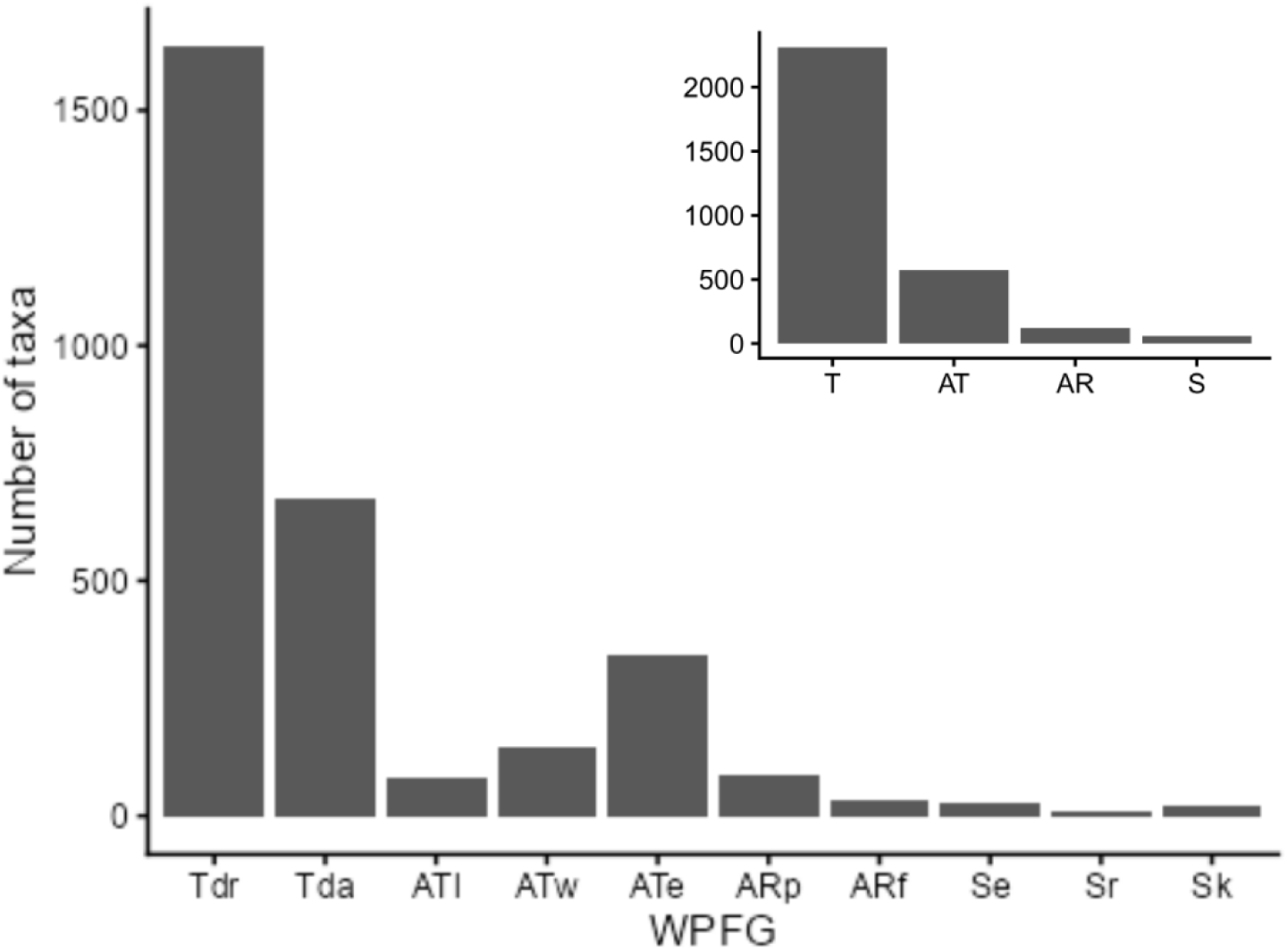
Distribution of species (or lower level) taxa among functional groups at two levels of the hierarchy. Main plot shows the count of species in each WPFG in the 10-category classification, while the inset shows the distribution when grouped by broader water tolerance, dropping the lowest level. WPFG in both plots are ordered from left to right in approximate order of increasing tolerance for permanent saturation or inundation.

### Inter-observer consensus

Of the total 3062 identified taxa at species-level or below, 1392 (45%) appeared in only a single list and could not be assessed for inter-observer variability with current information. Under the 10-category classification, the remaining taxa appeared in between 2 and 13 different lists, within which they were assigned to between 1 and 4 WPFG classes in different sources. Of 1670 taxa appearing in more than one list, 1093 (65%) were always assigned to the same WPFG.

The highest observed weighted pairwise cross-classifications between WPFG in the 10-category classification were at the dry end of hydrological niche space (Fig. 4a). Terrestrial dry - terrestrial damp confusion alone accounts for the largest proportion (41.5 %) of conflicted pairwise taxonomic assignment across all listings. Conflict between terrestrial damp (Tda) and amphibious emergent tolerators (ATe) was the second most common pairing, accounting for a further 13.8 %. No other combination of WPFG pairs exceeded 8 % of total observed cross-classifications (Fig. 4a). Prevalence-adjusted cross-classification ratios differed considerably (Fig. 4b), with submerged *r* and *k* selected (i.e., Sr – Sk), and amphibious plastic and floating responders (ARp – ARf) both being cross-classified > 15 times expectation given their prevalence. Amphibious plastic responders and low-growing amphibious tolerators (ARp–ATl) were also cross-classified more than 10 times as frequently as expected. In general, the greater the contrast in either life form (e.g., woody vs low-growing) or preferred hydrological conditions (e.g., T vs S) between WPFG pairs, the lower the prevalence of cross-classification.

**Figure 4.**
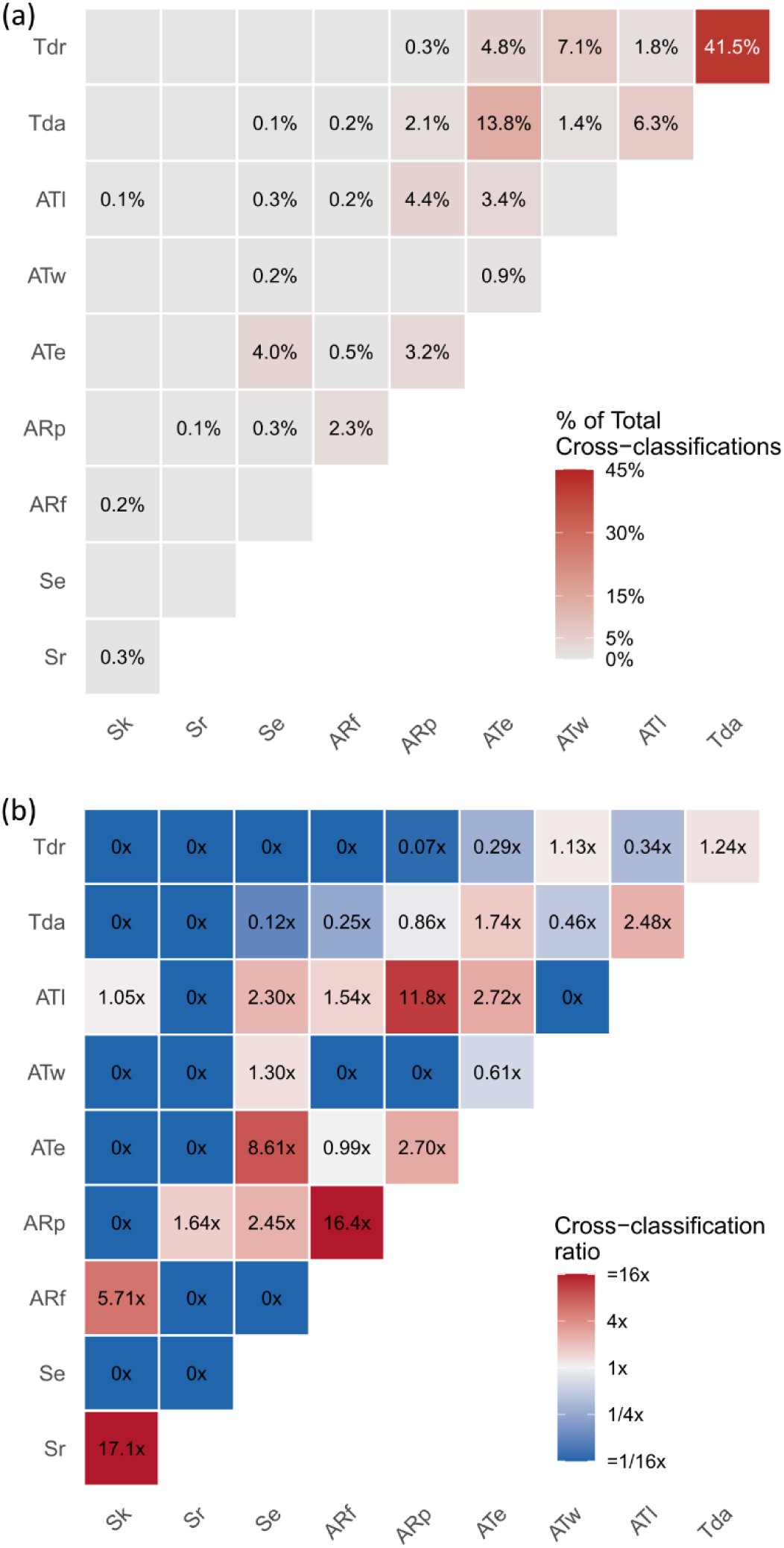
Heatmaps of cross-classification outcomes in pairwise comparisons of WPFG assignments made by different researchers under the 10-category classification. (a) Observed weighted pairwise cross-classification (b) Prevalence-adjusted cross-classification ratio (number of times observed cross-classifications exceeded expectation based on WPFG prevalences).

## Discussion

### Contribution of this database

This database provides two advantages that should advance trait-based analyses of environmental water requirements and trait–environment relationships for wetland plants. First, it increases transparency and reproducibility in assigning species to WPFG categories. It provides a central repository of the most comprehensive listing of individual taxa yet compiled, together with explicit information on the study systems from which each listing originates, plausible alternative WPFG classifications where taxa have previously been assigned differently, and a confidence score for assignment to a specific category. It also provides a centralised resource to which future lists and refinements can be added. As new study systems and taxa are incorporated into the WPFG framework, contributing researchers can update the database to a new version while retaining the digital identifier, enabling subsequent users to access the latest version.

### Caveats and limitations

Two main limitations associated with the use of WPFG are (i) uncertainty in assigning species to functional group categories and (ii) data gaps arising from the patchy spatial and taxonomic coverage of previous studies applying the method. These are considered separately below. Ideally future development of the framework would work toward increasingly associating WPFG categories with specific measurable traits (e.g., inducible vs constitutive aerenchyma) or with quantifiable characteristics and test its bioregional utility both within Australia and internationally.

#### Guidelines for species assignments

Although WPFGs essentially reflect variation in functional traits, many of the traits needed to confidently assign species to categories are available for relatively few species. In practice, assignments usually proceed through expert opinion, field observation, experimental trials or some combination of these approaches (Table 1). Confident assignment can require mesocosm or other experimental work that is time-consuming and expensive; consequently, field-survey evidence and expert knowledge are often used as practical substitutes. Expert knowledge can also be limited where little is known about the conditions under which seeds of a taxon germinate or establish. Ideally, diagnostic trait states could be identified, but this is currently of limited practical value because many potentially diagnostic traits—for example, inducible versus constitutive root aerenchyma to differentiate Tda and ATe—are not widely known for most species.

The most frequently observed cross-classifications (Fig. 4a) arise in differentiating terrestrial-dry from terrestrial-damp species and the latter from true amphibious species, especially emergent tolerators (ATe). The concentration of uncertainty around the Tdr–Tda–ATe boundary is ecologically plausible because these groups occupy the transition between predominantly terrestrial habitats and sites subject to periodic saturation or inundation. Species in this part of the hydrological gradient may appear terrestrial during dry phases but germinate, establish or persist under damp, waterlogged or shallowly inundated conditions. Seed-bank emergence studies show that functional-group composition can differ among rainfall, waterlogged and flooded treatments after an extended dry phase, and that seed-bank composition can vary with water depth and inundation zone (Kelleway et al. 2020, Ling et al. 2022, Ling et al. 2023). These examples suggest that classifications near the Tdr–Tda–ATe boundary should be interpreted with reference to evidence source, hydrological phase and study context rather than treated as fixed attributes in all settings. Guidelines for species assignments to WPFG categories are available (e.g., Brock and Casanova 1997, Casanova and Brock 2000, Casanova 2011), and for the seven-category classification are condensed here to a binary decision tree (Fig. 5).

**Figure 5.**
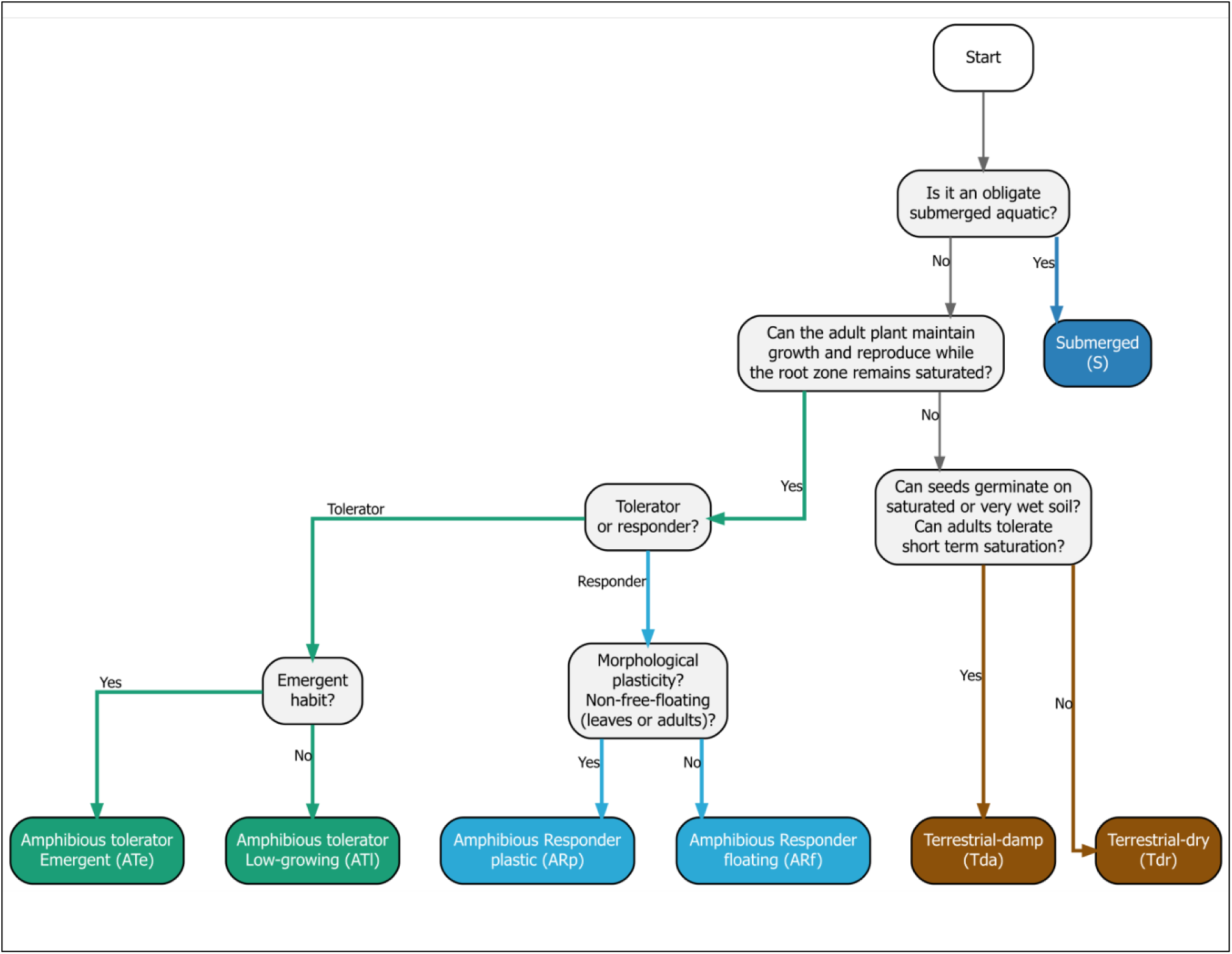
WPFG classification decision tree for the original seven-category classification. Decision points based on Brock & Casanova (1997) and Casanova & Brock (2000).

#### Bioregional gaps in WPFG application

At a national level, WPFG application in aquatic-plant research is uneven (Fig. 2). Tropical and subtropical east-central Australia, temperate south-western Australia and Tasmania have seen little or no prior application of WPFG in research. This pattern appears to be partly historical, reflecting the concentration of early WPFG research and collaboration in south-eastern Australia and the Murray–Darling Basin, as well as the availability of funding for monitoring and reporting environmental-water outcomes. The limited application in some regions may also reflect difficulty applying the classification where plant-trait data are scarce. Working at the level of species groups allows greater consistency of reporting and target setting, with clear advantages for developing a common management and reporting framework (Campbell et al. 2014). Whether the lack of published studies reflects limited research focus on trait-based functional responses outside the Murray–Darling Basin, poor suitability of the WPFG framework, limited trait data or another factor remains unknown. Given successful application of the WPFG framework in predicting plant-species responses in palustrine wetlands outside the Murray–Darling Basin (Campbell et al. 2014, Johns et al. 2015b, Deane et al. 2017, Deane et al. 2025), WPFG could be similarly useful in other regions. The collated database should help facilitate application to new bioregions and study systems. However, the database is likely to be most robust for south-eastern Australia and the Murray–Darling Basin, where most source studies were conducted. Applications in under-represented regions should therefore be locally validated, particularly where species occur under hydrological, climatic or wetland conditions not represented in the source lists.

### Prior uses of water plant functional groups

In contrast with the relatively restricted geographical focus toward south-eastern Australia, research applications of WPFG have been diverse. A common motivation is the increased generality of analysing at the level of WPFG, either for predictive or reporting purposes. Campbell et al. (2014) showed that analysis at this level can clearly differentiate contrasting taxonomic outcomes of wetlands as originating from common functional responses to flooding history. Similarly, Johns et al. (2015b) found WPFG more reliable at detecting community-level responses to water regime than growth forms or wetland indicator categories in palustrine wetlands. Deane et al. (2017) used WPFG to model vegetation responses to inundation in palustrine groundwater dependent wetlands, finding good predictive ability, with validated WPFG flooding-response curves used to predict potential equilibrium zonation under various groundwater drawdown scenarios (Deane et al. 2018). Deane et al. (2025) take a similar approach but given the independence of trait axes associated with flooding-tolerance and productivity (Pan et al. 2022), argue for the use of the more limited categories based on hydrology (T, A, S; or T, AT, AR, S) for predicting high-level impacts of hydrological change. Casanova (2011) exploited generality in response in the opposite direction, characterizing the water regime at different locations along an intermittent river floodplain based on knowledge of the water requirements of the WPFG of the observed vegetation.

WPFG has also been used in setting resource condition targets in the South Australian Murray River and associated floodplain (SA Murray River Corridor) as they provide a simplified transferrable approach in systems with large numbers of species. The Living Murray (TLM) vegetation condition targets for Lakes Alexandrina and Albert (Robinson 2014, Nicol et al. 2026a) and Ramsar Limits of Acceptable Change (Nicol 2016) largely used the Level 2 (four category; see Fig. 1) classification in Casanova (2011) to determine vegetation condition. Similarly, TLM understorey vegetation condition for the Chowilla Floodplain used a functional approach (Robinson 2014, Nicol et al. 2026b) based on Casanova (2011). The Chowilla Floodplain condition targets were adopted throughout the SA Murray River Corridor between the New South Wales border and Wellington, including the managed Pike and Katarapko floodplains (Environmental Watering Unit 2025). However, it is recognised that there are limitations at the dry end of the spectrum using the Casanova (2011) 10-category framework (see Fig 4) and an additional category that includes species intolerant of inundation but only found in areas periodically inundated has been added for target setting in floodplain environments.

WPFG has also supported soil seed-bank studies by providing a functional lens for interpreting regenerative capacity, drought resilience and restoration potential (Kelleway et al. 2020, Ling et al. 2022, Ling et al. 2023). In addition, WPFG can simplify environmental-impact monitoring and reporting where indicators must be readily communicated to managers and regulators, such as monitoring changes in wetland versus dryland vegetation cover.

Other studies have used the classification as a filter to separate obligate terrestrial and wetland plants before comparing diversity-scaling (alpha, beta) patterns (Deane et al. 2016) or separately analysed native and non-native cover within functional groups to partition impacts across hydrological niche space. This approach, using a hybrid three-category classification (Tdr, Tda, A) was used to develop a monitoring and reporting framework to help detect and communicate environmental impacts on threatened groundwater-dependent wetland plant communities from potential hydrological disturbance resulting from longwall mining (Brownstein et al. 2014, Johns et al. 2015a). Here, monitoring and reporting protocols used a before–after– control–impact (BACI) design to detect changes in WPFG cover indicative of drying, with significant differences in vegetation responses between control and mining-affected sites serving as a trigger for regulatory intervention. Deane et al. (in prep) compared native and non-native cover within the 10-category classification in agricultural and conservation landscapes to quantify relative impacts on native plant cover. Together, these applications illustrate the broad utility of WPFG for both predictive analysis and reporting across wetland research and management.

## Methods

Compilation of the database followed three broad stages (Fig. 6): (i) collation and harmonization of taxonomic nomenclature of source data, (ii) analysis and validation of inter-observer consensus and WPFG cross-classification, and (iii) final database consolidation. Each stage integrated integrity checks, ensuring domain validity (checking allowed WPFG categories) and no duplication of records. Each compilation stage involved is discussed below and the R code and raw data used to implement the analyses can be obtained from the github repository (https://github.com/deaned01/WPFG-database).

**Figure 6.**
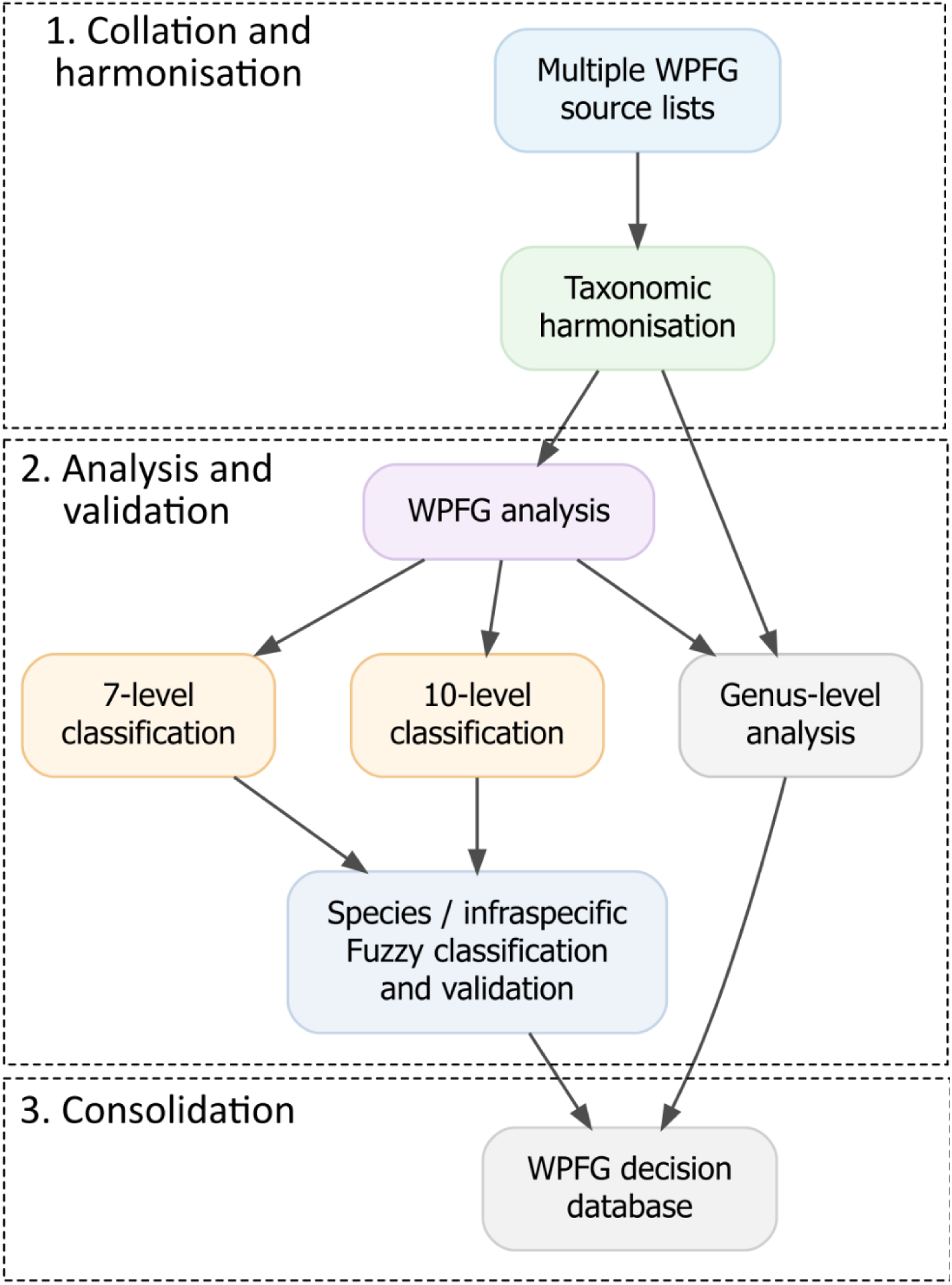
Key steps and phases in creating the Australian Water Plant Functional Group database.

### Stage 1 - Collation and taxonomic harmonisation

This database grew out of an informal collaboration between all co-authors initiated in 2024. Following two online workshops, all participants were asked to contribute lists of their own or to identify lists from published studies. Lists assigning species to water plant functional group categories (Table 1) were collated from (i) peer-reviewed primary sources (Brock and Casanova 1997, Casanova and Brock 2000, Leck and Brock 2000, Price et al. 2010, Casanova 2011, Campbell et al. 2014, Casanova and Powling 2014, Johns et al. 2015b, Deane et al. 2017, Bowen 2019, Ling et al. 2019, Mason et al. 2022), (ii) publicly available ‘grey’ literature, typically reporting on government agency monitoring programs (MDFRC and CSIRO 2014, Dyer et al. 2025), and (iii) personally curated lists created by experienced aquatic botanists working in Australian rivers and wetlands (M. Casanova, J. Nicol, K. Ward, P. Driver). Some of the later lists are extensions of existing published or shared list, for example the Flow-MER database was built on Bowen (2019). This knowledge means that a given taxon assigned to the same WPFG in two separate lists cannot be assumed to represent independent assignments. This dependence of records is accounted for in the compilation of the database (see ‘Reconciliation of classifications within source lists’).

Source data were collated within a single long format comma separated value file, with each record comprising the taxon name reported by the source, its assigned WPFG, a source identifier (e.g., first_author_year), and a chronological index representing the approximate order in which source lists were produced. The chronological index was an integer value, assigned according to the date the list was compiled, adopting either the date of any associated publication or the date of most recent additions to the list for personal communications. The integer coded chronological index was used to combine lists sequentially to create the database.

The collated WPFG lists were imported into R, where taxonomy was harmonized using R package APCalign (Wenk et al. 2024). This package facilitates aligning of vascular plant scientific names in Australia to the currently accepted name in the Australian Plant Census or Australian Plant Names Index. The taxonomic lookup retained the original and aligned names, APC accepted and suggested names, family, genus, scientific name, taxonomic rank and status, and information describing the basis for name alignment and taxonomic updating. A unique internal taxon identifier (taxon_id) was generated to link records resolving to the same taxonomic entity after aligning with accepted names. All original records were retained in the master database, including WPFG assignments made at genus or higher taxonomic ranks. However, higher-level assignments were not assumed to apply to individual species because congeners may differ functionally. Records resolved to species or infraspecific rank were therefore identified as eligible to provide evidence on species-level WPFG assignments, while higher-rank records were excluded from subsequent species-level classification. These were treated using a separate genus-level analysis to provide an indication on how well conserved WPFG classifications were within genera.

#### Reconciliation of classifications within source lists

After taxonomic harmonization, it was not unusual for two or more names within the same source list to resolve to the same taxon; their WPFG assignments could be identical or conflicting. To avoid treating taxonomic synonyms or duplicate source records as independent classifications, records were reduced to a single WPFG assignment for each unique taxon × source combination. To do this, we retained the same WPFG level if no conflict arose. If multiple classifications resulted in a clear numerical majority, the most common level was used. Where classifications were tied, a predefined WPFG hierarchy was used to select a single assignment, where increasing tolerance for greater duration and depth of flooding was preferred (i.e., Tdr < Tda < ATl < ATe < ARp < ARf < Se < Sr < Sk). All original names, classifications, number of contributing records and methods used to select the source-level classification were retained to provide an audit trail.

### Stage 2 – analysis of inter-observer consensus and validation

#### Seven- and ten-category WPFG systems

Owing to the change from the seven- to the ten-category WPFG classifications across datasets, we analysed taxon affinity in both. Because the seven-category system can be obtained by simplifying the ten-category system, all records could be analysed at this level. To do this, 10-category source classifications were converted to this common resolution, with Sr, Sk and Se re-coded as S and ATw re-coded as ATe. Only sources produced after expansion of the classification system around 2010 could be used for the ten-category analysis (Table 1).

#### Non-independence among historical classifications – analysis of assignment chains

An important feature of predominantly expert-opinion based classifications such as WPFG is establishing inter-observer agreement. We therefore undertook a range of analyses to provide guidance to users on the most reliable species-functional group assignments and to identify where most uncertainty lies in assigning species to different WPFG. We also calculated a fuzzy classification, which assigns a score to each species for each level of both the seven- and ten-category classification reflecting how the evidence is distributed over all assignments.

As mentioned above, a key challenge in comparing inter-observer consensus when collating this database was lack of explicit knowledge of whether species assignments from earlier lists were actively reviewed or automatically adopted as previously assigned. This distinction is important in establishing species affinities for a particular WPFG because careful inspection of a prior classification would offer an independent validation of the suitability of that class for that species. We assumed that in most cases citations were circular (adopted with limited individual review) and thus used a temporal weighting parameter (α = 0.2), meaning subsequent repeat classifications of a previously recorded taxon were incorporated as a partial independent assignment. This effectively assumes that 80% of new assignments naïvely accepted prior classification of species where one existed. We later tested for sensitivity of cross-classification metrics to this design decision.

The calculation itself proceeded as follows. For each taxon, source-level classifications were ordered chronologically to form an assignment chain. The first observed classification for each taxon received a weight of 1.0, indicating an independent assignment. Any subsequent unchanged classification was assigned a penalized weighting (denoted α) of 0.2, representing potentially inherited rather than independently reassessed evidence. Any change to a different WPFG in subsequent lists was interpreted as evidence of an active independent reassessment and received full weight (1.0). Thus, for example, if A and B denote two different WPFG categories, an assignment chain A → A → B received weights 1.0 + 0.2 + 1.0, whereas A → A → A received 1.0 + 0.2 + 0.2. The sum of dependence weighted assignment chain scores is referred to as ‘effective evidence’ and used in fuzzy classification and comparative weighting.

The original submerged category (S) was assigned additional treatment in the ten-category analysis because several lists included this level (essentially reflecting an 11-category classification), where S represents a coarser parent classification that offers some information toward its sub-categories (Sr, Sk and Se). Transitions between S and one of these sub-categories (e.g. S → Se or Se → S) were classified as hierarchical refinements rather than genuine disagreements and received the same weight as repeat observations (0.2). Only transitions between distinct S sub-categories (e.g. Se → Sk) were treated as genuine classification changes.

#### Fuzzy WPFG classification

To determine relative support for each taxon against each WPFG category, we summed the effective evidence for each WPFG within each taxon. Fuzzy membership was then calculated by dividing the effective evidence associated with each WPFG by the total effective evidence for that taxon. Fuzzy membership values therefore sum to one within each classification system. For the seven-category system, each source assignment contributed to one of the seven WPFGs, while in the ten-category system, explicit assignments contributed entirely to their corresponding WPFG, but an unresolved S assignment contributed equally to the weights for Sr, Sk and Se for that taxon.

The WPFG with the greatest fuzzy membership was reported as the ‘best guess’ for that taxon and classification resolution. The second-ranked classification and its support were also retained. Here, support represents the proportion of total dependence-adjusted evidence assigned to the best guess, while margin represents the difference between support for the first- and second-ranked classifications. Support describes the degree to which the available evidence favours the best guess, whereas the margin describes how clearly it is separated from its principal alternative.

#### Evidence depth

We differentiate the quantity of historical evidence underlying each classification (the number of source lists in which the taxon appears) from the consensus among classifications. We thus present the number of distinct source lists contributing species-level evidence (‘n_source’), and the ‘effective_evidence’ (the sum of dependence-adjusted source weights). This distinction was retained because complete agreement does not necessarily imply strong evidence. For example, taxa occurring in only one source list must have a support of 1.0 but will also have effective evidence of 1.0 (the minimum possible). In contrast, taxa classified consistently across multiple sources will have higher effective evidence. Conversely, taxa represented in many sources can still have lower support if repeated reassessments have resulted in disagreement. Support, evidence depth, the first-to-second classification margin, and the complete fuzzy memberships were retained in the final database as they provide more transparent descriptions of classification certainty and evidence availability.

#### Genus-level analysis

Because higher-level WPFG assignments cannot safely be automatically propagated to individual species, genus-level information was analysed separately. This offers both guidance on assigning taxa with only genera-level identification and a descriptive measure of conservation of genera-level functional-group affinity. This analysis did not contribute evidence to the species-level seven- or ten-category WPFG classifications.

For each genus, species-level fuzzy memberships were aggregated across classified species, with each species contributing equally regardless of the number of historical source lists in which it occurred. For WPFG (*k*) within genus (*g*), genus-level membership was calculated as the arithmetic mean of the corresponding species-level fuzzy memberships:

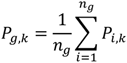

where (*n_g_*) is the number of classified species in genus (*g*), and *P_i,k_* is the species-level fuzzy membership of species, *i* for WPFG, *k*.

Equal weighting of species was used to describe conservation of WPFG category for the species within a genus. That is, a species represented by many source lists did not receive greater weight in the genus-level summary than a species represented by one source. Analogous to the species-level assignments, for each genus, the dominant WPFG, its genus-level membership, the second-most supported WPFG, the margin between them, the number of WPFGs represented, and the complete genus-level fuzzy membership distribution were retained. Note that historical WPFG assignments explicitly made at genus or higher taxonomic rank were also retained but were treated as descriptive observations of previous classification practice, not as evidence for assigning those WPFGs to individual species. Genus-level information explores how conserved WPFG classifications appear to be among species within a genus, while also providing guidance on the most likely WPFG and alternative choices at this level.

#### Cross-classification among WPFGs

Observed inter-observer transitions between WPFGs were collated to identify pairs of functional groups most likely to be cross-classified. Only genuine transitions between non-overlapping classifications were treated as cross-classification events. Repeated classifications were excluded, as were hierarchical transitions between S and Sr, Sk or Se. Cross-classification was calculated from transitions between successive source assignments using a symmetric analysis (pooling cross-classifications in either direction) of reciprocal transitions between pairs of WPFGs. Cross-classification events were weighted using the same dependence framework as the classification analysis and therefore describe the relative distribution of dependence-adjusted historical disagreements among WPFG categories. Two different measures were calculated, an observed proportion of cross-classification and one comparing the observed proportion to an expectation conditional on the prevalence of each WPFG.

##### Cross-classification approach

Pairwise cross-classification was evaluated separately for the 7-category and 10-category classifications, using the weighted effective evidence. Reciprocal transitions between the same two WPFGs were pooled, so that A-to-B and B-to-A contributed to the same unordered pair. The weighted observed cross-classification (presented as a heatmap; Fig. 4a) for each pair represents its proportion of all observed cross-classification weight. For pair i,j, the observed proportion was calculated as:

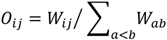

where Wᵢⱼ is the total dependence-adjusted weight for cross-classifications between WPFGs i and j, and the denominator is the total cross-classification weight across all unordered pairs. Thus, the observed cross-classification values sum to 1 (100%).

##### Prevalence-adjusted pairwise ratio

Because common WPFGs have more opportunities to be cross-classified, observed pairwise proportions were also compared with the frequencies expected from the marginal prevalence of each WPFG. Let *p*ᵢ denote the proportion of weighted assignment evidence belonging to WPFG *i*. Under an independence model, the expected probability of an unordered pair i,j is:

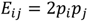

The analysis conditions this expectation on a mismatch (i.e. two different WPFGs), giving:

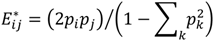

Here, *i* and *j* identify the focal WPFG pair, while *k* indexes all WPFG categories in the relevant classification system (7 or 10 category). The denominator is the expected probability that two independently drawn weighted assignments belong to different WPFGs. The prevalence-adjusted cross-classification ratio, *R*, was then calculated as:

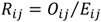

A ratio of 1 indicates cross-classification at the frequency expected from WPFG prevalence; values >1 indicate more cross-classification than expected, and values <1 indicate less. This metric does not identify which WPFG is more correct but rather describes whether a pair is cross-classified disproportionately often or seldom, after accounting for how common the two WPFGs are. Observed cross-classification and the ratio were interpreted alongside the number of cross-classification events and distinct taxa contributing to each pair, because large ratios can occur for rare pairs with small evidence bases. Both cross-classification results are presented in pairwise heatmaps (Fig. 4) for the 10-category classification (7-category results were qualitatively identical allowing for the lesser number of categories).

##### Interpretation across classification resolutions

The 7-category and 10-category systems were compared descriptively using the same definitions and, for the heatmaps, the same scale where appropriate. Similar pairwise patterns across resolutions indicate stability of historical cross-classification structure, whereas newly prominent or disproportionately cross-classified pairs in the 10-category system may identify distinctions that have been applied less consistently. These analyses assess historical classification consistency and do not establish biological correctness or superiority of either classification resolution.

#### Validation and sensitivity analysis

To test the rigour of the classification framework, we used structural checks, synthetic assignment chains, empirical diagnostics and sensitivity analyses. Structural validation confirmed that fuzzy memberships summed to one, only valid WPFG categories occurred within each classification system, and each taxon contributed no more than one classification per source. Synthetic chronological chains were used to verify the dependence-weighting rules. For example, A → A → A produced effective evidence of 1.4 (under α = 0.2) and A → B produced 2.0. Hierarchical evidence handling was separately checked to ensure that transitions between S and Sr, Sk or Se were identified as refinements rather than genuine conflicts. Ten-category fuzzy calculations were additionally checked to ensure that S evidence was distributed among Sr, Sk and Se as specified. Sensitivity to the assumed dependence penalty was assessed by recalculating classifications across α values of 0, 0.1, 0.2, 0.3, 0.5 and 1.0. These span assumptions ranging from no additional evidence being attributed to an unchanged repeated classification (α = 0) to treating all repeated classifications as fully independent (α = 1). Best-guess classifications remained unchanged across the full range of α for 94.7% and 94.6% of taxa in the seven- and ten-category analyses respectively, indicating that the selected WPFG was generally robust to the assumed dependence penalty.

### Stage 3 - Final database outputs

The final user-facing database reports taxonomic identity, including APC family and genus, together with separate seven- and ten-category advice on the most likely WPFG. For each classification it reports the best guess WPFG, support for the best guess, strongest alternative classification, support for that alternative, the margin between first-to-second best, the number of contributing sources in which the taxon appeared, the effective evidence (weighted by dependence), and the complete fuzzy WPFG memberships. The separate genus summary reports species-derived genus-level fuzzy composition and historical genus-level WPFG assignments. This information is explicitly descriptive and does not contribute to species-level recommendations. The database also retains the original source names and taxonomic provenance to facilitate matching of new species lists and reproducibility of database releases. Each release records the database version, source input file, and APC taxonomic resource/version used for name harmonisation.

#### Use of generative artificial intelligence

ChatGPT (OpenAI; GPT-5.6 Sol) was used during development of the analytical pipeline. It was used to assist in R code development and debugging, implementation and validation of the fuzzy-classification framework, development of the reproducible database workflow, and in confirming methodological documentation. All analytical decisions, code execution, validation, interpretation of results and final manuscript content were undertaken and reviewed by the authors.

## Acknowledgements

We acknowledge the contribution of all researchers to the development of the Water Plant Functional Groups classification and its use in research, particularly Prof. M. Brock. DCD was supported by a grant from the Australian Research Council (grant # DE240100477). FJD, AT and TJM were supported by The Commonwealth Environmental Water Holder Flow-Monitoring, Evaluation and Research (Flow-MER) Program. Keith Ward, Daryl Nielsen and Patrick Driver contributed data to the Australia-wide database held by Michelle Casanova; Max Finlayson, Arnold van der Valk, Cassie James, Sam Capon, Adrian Clements, Ray Froend, Robert Godfree and Jane Roberts contributed to discussion workshops held under the aegis of the Australian Centre for Ecological Analysis and Synthesis between 2012 and 2014.

## Author contributions

DCD initiated the collaboration, collated the database, performed all inter-observer analyses and wrote the first draft of the paper. All authors contributed to the development of the database through participation in online workshops. All authors contributed to the writing and revision of the draft and approved the final submission. Order of authorship was determined alphabetically except for first and last authors.

## Data availability statement

The Water Plant Functional Group Database Version 1, including the user database and supporting analytical and validation files, is publicly available through the Zenodo repository (https://doi.org/10.5281/zenodo.23128087). The source data, R code and reproducible computational environment used to generate the database are available through GitHub (https://github.com/deaned01/WPFG-database), with the corresponding workflow archived as release v1.0.0 (https://github.com/deaned01/WPFG-database/releases/tag/v1.0.0).

